# Exposure to neodymium blunts the hypoxic ventilatory response in fathead minnows (*Pimephales promelas*)

**DOI:** 10.1101/2024.11.29.626036

**Authors:** Natalie M. Nykamp, James C. McGeer, Erin M. Leonard

## Abstract

In hypoxia, the initial response in vertebrates is hyperventilation, known as the Hypoxic Ventilatory Response (HVR), which is a physiological reflex that allows fish to maintain adequate oxygen uptake. The severity of hypoxia in aquatic ecosystems is growing due to anthropogenic impacts. This is a concern with the recent evidence that metals can affect the ability of fishes to mount the HVR, potentially impacting survival. As Rare Earth Elements (REEs) increase in demand with the shift to a low-carbon economy, there is a critical need to understand their environmental consequences. Neodymium (Nd) is used in green technology and is one of the most critical REEs. Here, we investigate whether exposure to Nd will blunt the HVR in a toxicological model, the fathead minnow (*Pimephales promelas*). The fathead minnow will be exposed to hypoxia and Nd, and the ventilation rate will be observed and compared to controls to determine if there is a blunt in the HVR when the fish are exposed to both hypoxia and Nd. Nd caused a 31% decrease in the HVR and Nd accumulation in the gills was below the detection limit (LOD: 0.0681 µg/L). Toxicity testing with REEs during this time of economic growth is imperative for the protection of aquatic life in Canada.

## Introduction

Low-oxygen environments are naturally occurring in aquatic ecosystems and can be heightened by anthropogenic activities such as nutrient loading, agriculture, and urbanization (see Tellier et al., 2022 for review). Hypoxia is described as zones in which the dissolved oxygen (O_2_) concentration falls below 2 mg O_2_/L. The severity and incidence of hypoxia is greatly increasing with eutrophication due to nitrogen and phosphorus runoff. Increasing temperatures due to climate change may exacerbate the impacts of hypoxia (Friedrich et al., 2014), creating an urgency in understanding the impacts of hypoxia on fish. Fish have adaptations to tolerate these low oxygen environments using a number of physiological processes such as aquatic surface respiration (ASR), bradycardia, behavioral changes, and the hypoxic ventilatory response (HVR) (Abdallah et al., 2015; Perry et al., 2009; Small et al., 2014). These processes are mainly initiated by the activation of the neuroepithelial cells (NECs) in gill filaments, which are responsible for sensing hypoxia and sending information to the central nervous system (see Zachar and Jonz, 2012 for review). The HVR is arguably one of the most important responses to hypoxia, present in many animal models, and describes the increase in ventilation as an immediate response to hypoxia (see Milsom, 2012 for review). This increase can be seen as an increase in the frequency of breaths and/or increase in the amplitude of breaths (Mandic et al., 2018), helping organisms maintain homeostasis and meet metabolic demands.

As of 2023, the price of rare earth elements (REEs) can be linked to economic development and geopolitical risks, differing them from other commodities due to their global applications and current dominance in markets and key industries (Liu et al., 2023). Canada, China, Australia, the United States, Brazil, and Russia all hold substantial deposits of REEs, with China being the leader in production, exportation, and consumption, producing over half of REEs (Government of Canada, 2023; Cuadros-Muñoz et al., 2024; Liu et al., 2023). There are significant REE deposits in Northern Canada (Government of Canada, 2024) and in 2023, Canada released its Critical Mineral Strategy; a document outlining a list of minerals that is projected to be mined and supplied by Canada to major global markets. This list includes REEs due to their applications in permanent magnets for electricity generators and motors (Government of Canada, 2023). Neodymium-iron-boron magnets and NiMH batteries both containing REE’s are essential for electrical vehicles, green technologies such as wind turbines, and everyday items such as computers and phones (Jin et al., 2018). Due to the predicted increase in electrical vehicles, increased mining of REEs is expected globally over the next few decades, leading to increases in REEs found in aquatic environments (Vukov et al., 2016). However, essential information relevant to the environmental consequences of the mining of REE is deficient. Regions such as Canada, United States, European Union, and Australia are in the process of developing guidelines or criteria to protect aquatic life pertaining to REE’s due to the lack of REE toxicity data (Markich et al., 2024) and Nd. Currently, there are no Federal Water Quality Guidelines for the Protection of Aquatic Life for REEs in Canada (CCME, 2024).

Exposure to metals can impact the ability of a fish to respond to hypoxic environments (Baker et al., 2016; Fitzgerald et al., 2019). It has been previously demonstrated that exposure to copper (Cu) can affect the HVR in killifish (*Fundulus heteroclitus*), and three-spined sticklebacks (*Gasterosteus aculeatus*) (Baker et al., 2016; Fitzgerald et al., 2019). 100 µg/L Cu showed a blunt in the HVR in killifish, potentially causing a decrease in oxygen uptake in fish co-exposed to Cu and hypoxia (Baker et al., 2016). Additionally, lower concentrations (20 µg/L) of Cu led to elevations in ventilation rate of three-spined sticklebacks, which increased respiratory stress and metabolic costs (Fitzgerald et al., 2016).

The hypothesis tested in this study is that Nd will blunt the HVR in fathead minnows (*Pimephales promelas*). We predict that Nd will act in a similar manner as Cu, reducing the ability of fathead minnows to mount the HVR to hypoxia. Additionally, we explored whether inhibition could be explained by increases in Nd accumulation in the gills.

## Methods and Materials

### Animal husbandry

Adult fathead minnows (1.4 ± 0.10 g), *Pimephales promelas*, of both sexes were kindly supplied by Drs. Deborah MacLatchy and Andrea Lister (Wilfrid Laurier University) under the animal use protocol (AUP) R23011. Minnows were housed in tanks filled with aerated water at approximately 20°C (Fisher Scientific Traceable Excursion Trac Datalogging thermometer, Massachusetts, USA). Water chemistry parameters were monitored every 24-h with an average conductivity of 1120 ± 12.4 µs/cs and average pH of 8.3 ± 0.026 (YSI ProQuatro Handheld Multiparameter Instrument Ohio, USA). Protocols for animal care were approved by the Laurier’s Animal Research Ethics Board (AUP #R23008).

### Neodymium exposure

Tanks were placed in a water bath, which was equipped with two submersible water heaters set to 25°C and a circulating pump (EHEIM GmbH & Co. KG., Deizisau, Germany). Minnows were exposed to Nd in an acute 48-h exposure. Prior to the addition of fish, control and Nd-prepared tanks were prepared and allowed to equilibrate for 24-h. (dissolved concentration 92.55 ± 7.930 µg/L Nd). Two fish were added to each of the control and Nd tanks to undergo a 48-h acute Nd exposure. Over the period of 48-h the fish were exposed to100% O_2_ (air saturation) and food was withheld to mitigate any effects of digestion. Following which, another four tanks were prepared for 24-h for conducting ventilatory testing and placed in the water bath. This created 4 distinct testing scenarios (control normoxia, control hypoxia, Nd normoxia, Nd hypoxia; see Fig. 1).

**Figure 1.**
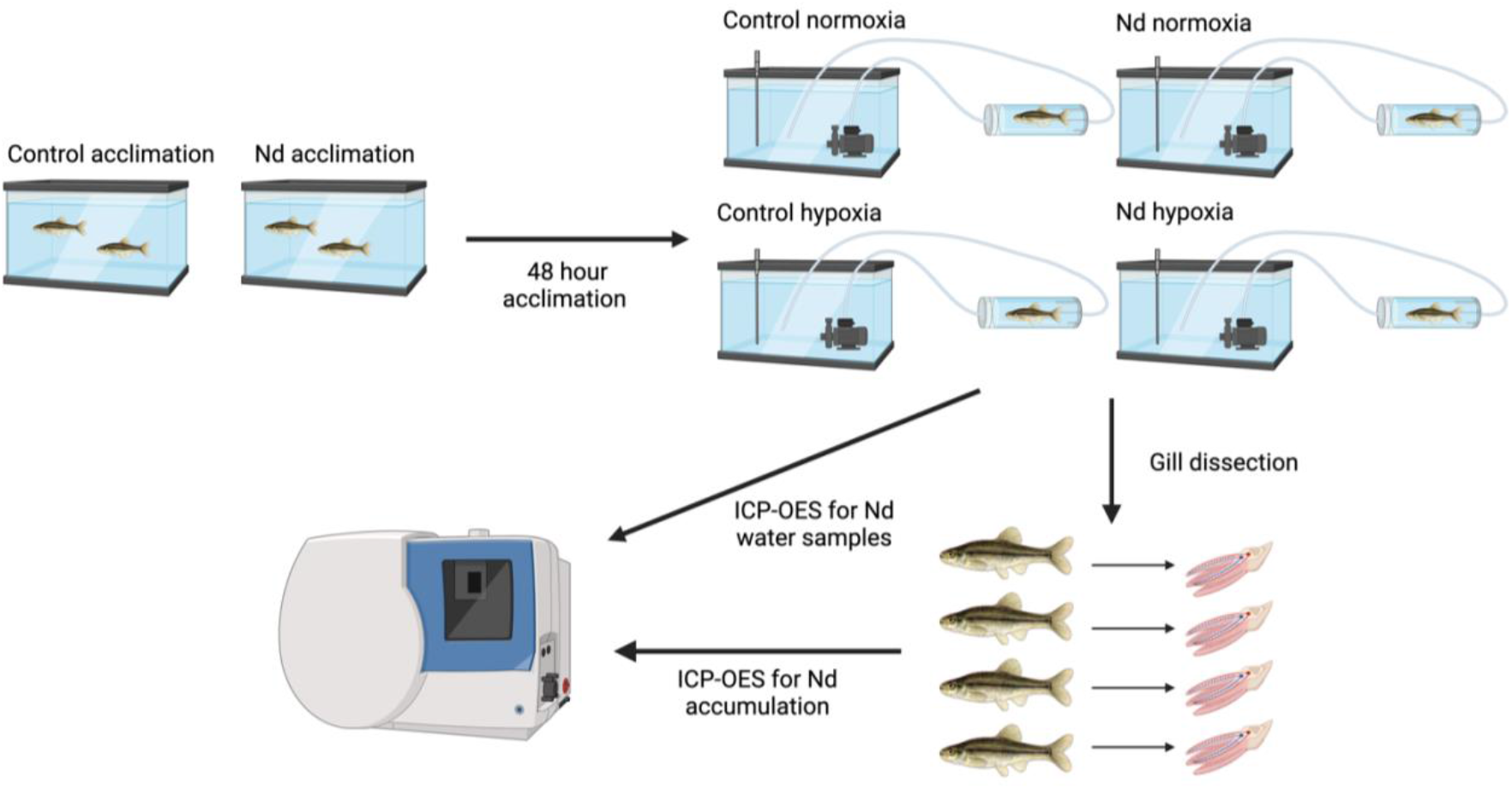
Overview of methods – The fish are acclimated for 48-h to either control water or the Nd exposure (92.55 ± 7.930 µg/L Nd). After the 48-h acclimation, each fish is placed in an individual fish chamber and are exposed to (A) control normoxia (100% O_2_), (B) control hypoxia (40% O_2_), (C) Nd normoxia (100% O_2_), or (D) Nd hypoxia (40% O_2_). A video of the fish is taken and analyzed to get a count of breaths per minute. The gills are dissected and ran through the ICP-OES to determine Nd accumulation in the gills. Water samples are also run through the ICP-OES to determine the Nd concentration in the water. (Created in BioRender. (2024) https://BioRender.com/w39e829).

### Ventilation rate measurements

The set-up consisted of individual fish chambers connected to a recirculating water system containing an EHEIM universal aquarium pump (EHEIM GmbH & Co. KG., Deizisau, Germany) and a microfiber optic oxygen meter sensor connected to a OXY4 multichannel (PreSens Precision Sensing GmbH, Regensburg, Germany) that was attached downstream of the individual fish chamber, used to monitor the DO concentration of each tank. The pumps were placed in one of the four tanks prepared for testing. The optic oxygen meter sensors were calibrated immediately before testing, using 100% O_2_ and 100% N_2_ and were placed in each of the four experimental tanks. Fish were placed in the individual fish chambers attached to recirculating systems in 100% O_2_ for 1-h to acclimate, as cortisol levels are known to drop to resting levels 60 minutes after handling (Philippe et al., 2023). After the 1-h acclimation period, the recirculating water systems were changed from 100% O_2_ to an exposure of either (A) control normoxia (100% O_2_), (B) control hypoxia (40% O_2_), (C) Nd normoxia (100% O_2_), or (D) Nd hypoxia (40% O_2_) (Fig. 1). The mix of O_2_ and N_2_ gas for the hypoxia exposure was controlled using a gas flow meter (Key Instruments, Pennsylvania, USA) and was adjusted to ∼ 40% dissolved oxygen (DO) over a 1-h period. A 10-min video recording of the individual fish chambers was then taken with an iPhone 11 camera (Apple, Zhengzhou, China). The video was analyzed to count the breaths of the fish, which were counted in 10 s intervals, three times throughout the video. These three counts were normalized to breaths per minute (bpm), and averaged for each fish. The DO% was recorded for each fish chamber. The fish were then taken out of the chambers and euthanized with a blow to the head followed by the severance of the spinal cord. Fish were weighed on an analytical scale, sexed, and the 8 gill arches of each fish were dissected and placed in pre-weighed Eppendorf tubes (Eppendorf, Hamburg, Germany) (Fig. 1).

#### Water and Gill Nd measurements

Dry weights of the gills (4.2 ± 0.49 mg) were determined after 24-h in an oven at 60ºC. Fish gills were digested using 1N trace metal grade nitric acid (Fisher Scientific, Ottawa, Canada) in Eppendorf tubes. Each tube was vortexed and placed back in the 60ºC oven for 48-h. Tubes were vortexed again and then placed in the centrifuge at 13,300 rpm for 15 minutes at 21ºC. Water samples were taken from the control and Nd acclimation tanks in 15 mL Falcon tubes (Fisher Scientific, Ottawa, Ontario, Canada) at 0-h, 24-h, and 48-h to test for total and dissolved Nd concentration (Table 1). Dissolved samples were measured by passing the water through a 0.45 μm filter (Fisher Scientific, Dublin, Ireland). All samples were acidified by adding approx. 0.05 mL of metal grade nitric acid. All metal concentrations in each water sample were measured using the Perkin Elmer Optima 8000 Inductively Coupled Plasma Optical Emission Spectrometer (PerkinElmer, Connecticut, USA). The standards used ranged from 7 to 1000 µg Nd/L (Inorganic Ventures, Christiansburg, Virginia, USA). Nd concentrations were tested using the Perkin Elmer Optima 8000 ICP-OES (Perkin Elmer, Massachusetts, USA; Table 1) and ion concentrations were tested using the Perkin Elmer Pinaacle 900T AAS Flame (Perkin Elmer, Massachusetts, USA; Table 1).

**Table 1:** Water Chemistry of Control and Nd Water Samples over 48-h Water chemistry parameters including Nd using the ICP-OES and ion concentration using the Flame Atomic Absorption Spectrometry, and pH and conductivity of the test water.

| Treatment | Nd<sub>dissolved</sub> ($\mu\text{g/L}$) | Na (mg/L) | Mg (mg/L) | K (mg/L) | Ca (mg/L) | pH | Conductivity ($\mu\text{s/cs}$) |
| --- | --- | --- | --- | --- | --- | --- | --- |
| Control | $-3.102 \pm 0.1586$ | $142.2 \pm 14.71$ | $36.66 \pm 1.745$ | $2.406 \pm 0.1058$ | $85.60 \pm 8.2993$ | $8.193 \pm 0.01900$ | $1089 \pm 21.76$ |
| Nd | $92.55 \pm 7.930$ | $142.2 \pm 12.13$ | $38.02 \pm 1.483$ | $2.422 \pm 0.1142$ | $99.96 \pm 7.743$ | $8.363 \pm 0.03530$ | $1148 \pm 11.12$ |

#### Statistical Analysis

Statistical analysis was performed using Prism 10.1.1 GraphPad (GraphPad Software, Boston, Massachusetts, USA). Two-way analysis of variance (ANOVA) was used to assess comparisons against multiple treatment groups, followed by Tukey’s Honest Significant Difference test to compare between specific pairs of group means. Statistical significance was allotted to differences with p < 0.05. Data has been reported as means ± S.E.M. (*N*).

## Results

### Ventilation Rates

Non-metal exposed *P. promelas* significantly increased their ventilation rate by 65.3% from normoxia (118 ± 4.29 breaths per minute (bpm)) to hypoxia (195 ± 12.5 bpm); Two-way ANOVA; p < 0.0001; Fig. 2. However, the increase in ventilation in Nd was significantly lower with an increase in ventilation rate of 37.8% from Nd normoxia (98.0 ± 2.53 bpm) to Nd hypoxia (135 ± 5.53 bpm); Two-way ANOVA; p < 0.01; Fig. 2. *P. promelas* significantly decreased their ventilation rate by 38.8% when comparing Nd hypoxia (135 ± 5.53 bpm) to control hypoxia (195 ± 12.5 bpm); Two-way ANOVA; p < 0.0001; Fig. 2. There was no significant difference between control normoxia (118 ± 4.29 bpm) and Nd normoxia (98.0 ± 2.53 bpm); Two-way ANOVA; p > 0.05; Fig. 2.

**Figure 2.**
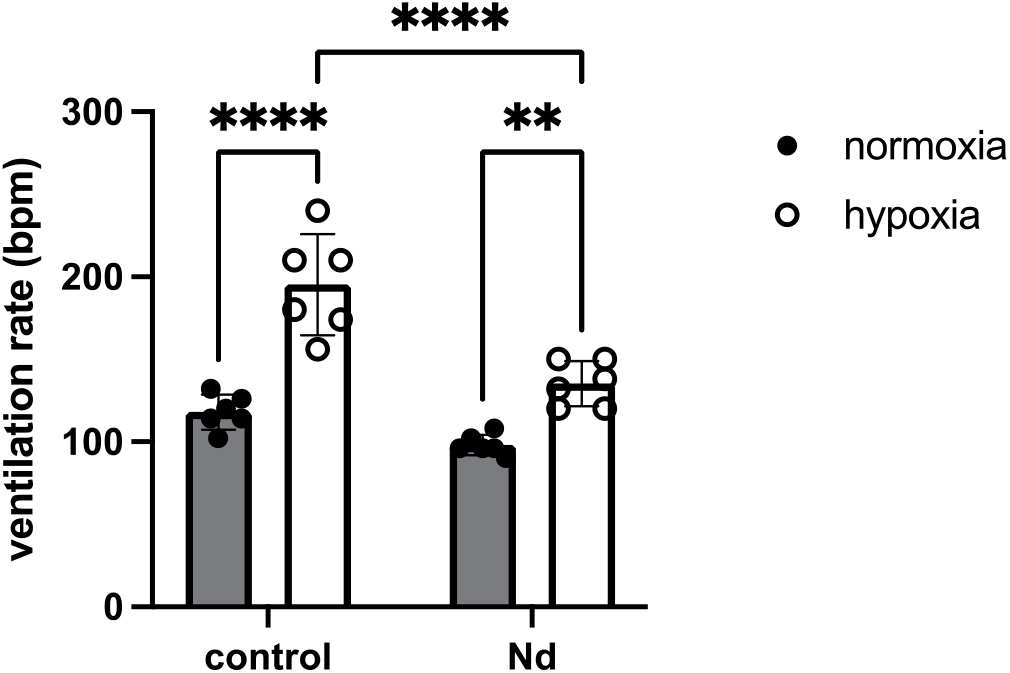
The effect of hypoxia and Nd on the ventilation rate of *P. promelas*. Mean ventilation rates (bpm) of fathead minnows exposed to control normoxia (black; n=6), control hypoxia (white; n=6), Nd normoxia (black; n=6), and Nd hypoxia (white; n=6). Nd dissolved exposure (92.55 ± 7.930 µg/L Nd) caused a significant blunt in the ventilation rate when comparing the control and Nd hypoxia groups (Two-way ANOVA; p < 0.0001). Asterix denotes significant differences (**p < 0.01, ****p < 0.0001).

### Bioaccumulation

Gill bioaccumulation did not exceed the detection limit of the ICP-OES (LOD: 0.0681 µg/L) in all 24 gill samples measured.

## Discussion

While disruptions to oxygen regulation and the HVR have previously been determined with divalent metals (Baker et al., 2016; Fitzgerald et al., 2019), this has not been tested with trivalent metals (such as Nd). This is a concern, given the increased demand due to their use in clean and green technologies. We originally hypothesized that trivalent REEs such as Nd would blunt the hyperventilatory response of *P. promelas* to hypoxia, and indeed, *P. promelas* exposed to the combination of hypoxia and Nd had a blunted HVR in comparison to controls. This significant blunt in ventilation frequency is observed in non-detectable Nd bioaccumulation levels, demonstrating significant physiological effects below bioaccumulation levels and a sensitive endpoint. With REE becoming economically important on a global scale with mines operating internationally, this study highlights the importance of toxicity testing concerning REE and sublethal endpoints for the protection of aquatic life.

### Water chemistry

These fish were exposed to Nd (92.55 ± 7.930 µg/L Nd) in water with high water hardness (∼250 mg/L as CaCO3) and a relative high pH (∼8.3); conditions that are considered protective if we are assuming that the premise of the Biotic Ligand Model (BLM) will work for Nd as has been shown for aluminum (Al^3+^) (Santore et al., 2017). The BLM predicts the toxicity of metals to aquatic organisms and works on the basis that dissolved metal concentrations are the bioavailable form of the metal that interacts with defined sites (i.e. biotic ligands) eliciting toxicity (see Paquin et al., 2002 for review). With non-detectable bioaccumulation of Nd causing whole organism level effects in *P. promelas* in protective water, it is assumed that less protective water would cause further impact on the organism. This study emphasizes the significance of understanding the interaction between metals and hypoxia. As there is little information on REE toxicity (Nd), there is a dependence on aluminum toxicity information. It is well established that Al toxicity is strongly dependent on factors such as pH, dissolved organic carbon (DOC), hardness, and temperature (Santore et al., 2017). Therefore, a typical water hardness equation may not be sufficient. Although this study does not assess the effects of water chemistry parameters on Nd bioaccumulation and incipient toxicity, the premise of the BLM is that metal toxicity is linked to metal bioaccumulation in the gills (Playle, 1998), which has also been shown with Al where increased mortality is correlated with increased Al bound to the gill (Santore et al., 2017). However, in the current study, there was non-detectable Nd bioaccumulation associated with a decrease in the ability of the fish to mount a full HVR, a sublethal endpoint which may greatly impact ecological success of the organism in the wild.

### Nd exposure caused a blunt in the HVR

With aquatic hypoxic zones increasing due to climate change and increased anthropogenic impacts on aquatic ecosystems (see Tellier et al., 2022 for review), a blunt in the HVR could impair increased O_2_ uptake to meet metabolic demands, potentially reducing survivability (Fig. 2; Abdallah et al., 2015; Small et al., 2014). Similar consequences have been shown in three-spined sticklebacks (*Gasterosteus acueatus*) exposed to Cu (20 µg/L) where hypoxia tolerance (reduction in Pcrit) was suppressed when combined with exposure to Cu (Fitzgerald et al., 2019). The inability to reduce Pcrit in Cu and hypoxia suggests that the fish may struggle to maintain aerobic metabolism. HIF proteins play a crucial role in regulating hypoxia responses as they modulate changes in Pcrit and can decrease a fish’s Pcrit in hypoxia improving hypoxia tolerance (see Prabhakar and Semenza, 2012 for review). It has been shown that a knockout of the HIF protein decreases the ability of zebrafish (*Danio rerio*) to withstand hypoxia for longer periods (Mandic et al., 2020). Due to the inability of the fish to reduce Pcrit when exposed to copper (Fitzgerald et al., 2019), this may suggest that Cu could be interacting with HIF proteins, reducing aerobic metabolism, but this remains to be empirically tested.

The mode of impact of Nd may be driven at the cellular level. For example, exposure of killifish (*Fundulus heteroclitus*) to Cu blunts the HVR, a response seen at the cellular level in response to a combined exposure of hypoxia and Cu (Baker et al., 2016). In addition, in cultured rat dorsal root ganglion neurons, exposure to Al^3+^ blocked voltage-gated Ca^2+^ channels with little to no recovery (Busselberg et al., 1994). Since Nd and Al both form 3^+^ cations in their natural states, this may demonstrate analogous chemical behavior and physiological effects. It is possible that Nd can interfere with voltage-gated Ca^2+^ channels, eliciting the blunt in the hyperventilation response. On the other hand, Al is a known neurotoxin in aquatic vertebrates, causing oxidative stress by accumulating in the nervous system and interfering with oxygen regulation (see Closset et al., 2021 for review). If Nd is functioning analogously to Al, then oxidative stress could mean an increase in reactive oxygen species (ROS) which can build up and affect the hypoxia-inducible factor 1 (HIF-1),which is a transcription factor that plays a crucial role in the HVR. Follow up investigations targeting the Ca^+2^ channels or disruptions to HIF-1 pathway may be warranted to discover the mechanism underpinning Nd mediated blunting of the HVR.

### Perspective of the research

Results from this study indicate that exposure of *P. promelas* to 92.55 ± 7.930 µg/L Nd_dissolved_ causes a significant decrease in the HVR, affecting the ability to hyperventilate when exposed to hypoxia and Nd. The combination of these two stressors could have impacts on the fish’s ability to live in hypoxic zones, as well as their ability to behave normally or meet metabolic demands, due to the possible decrease in oxygen uptake. This can affect their ecological success because decreased oxygen can affect metabolic rate, growth, feeding, and reproduction (Neubauer and Andersen, 2019). Understanding the implications of heightened anthropogenic inputs such as Nd, in combination with hypoxia, is imperative for the protection of aquatic life in freshwater ecosystems.

## Acknowledgements

The authors would like to thank the lab technicians at the Centre for Cold Regions and Water Science, Gena Braun and David McAlpine. The authors also thank Drs. Deborah MacLatchy and Andrea Lister for the fathead minnows. We would like to acknowledge funding from the Natural Sciences and Engineering Research Council of Canada Grants to EML (RGPIN-2023-05466).

## References

Abdallah, S. J., Thomas, B. S., & Jonz, M. G. (2015). Aquatic surface respiration and swimming behaviour in adult and developing zebrafish exposed to hypoxia. Journal of Experimental Biology. 10.1242/jeb.116343

Baker, S. (2016). Effects of copper on the acute ventilatory drive of killifish, Fundulus heteroclitus (dissertation).

Busselberg, D., Platt, B., Michael, D., Carpenter, D. O., & Haas, H. L. (1994). Mammalian voltage-activated calcium channel currents are blocked by PB2+, zn2+, and al3+. Journal of Neurophysiology, 71(4), 1491–1497. 10.1152/jn.1994.71.4.1491

Canadian Council of Ministers of the Environment. (n.d.). Canadian water Quality Guidelines for the Protection of Aquatic Life. CCME. https://ccme.ca/en/resources/water-aquatic-life

Closset, M., Cailliau, K., Slaby, S., & Marin, M. (2021). Effects of aluminium contamination on the nervous system of Freshwater Aquatic Vertebrates: A Review. International Journal of Molecular Sciences, 23(1), 31. 10.3390/ijms23010031

Cuadros-Muñoz, J.-R., Jimber-del-Río, J.-A., Sorhegui-Ortega, R., Zea-De la Torre, M., & Vergara-Romero, A. (2024). Contribution of rare earth elements is key to the economy of the future. Land, 13(8), 1220. 10.3390/land13081220

Fitzgerald, J. A., Urbina, M. G., Rogers, N. J., Bury, N. R., Katsiadaki, I., Wilson, R. W., & Santos, E. M. (2019). Sublethal exposure to copper supresses the ability to acclimate to hypoxia in a model fish species. Aquatic Toxicology, 217, 105325. 10.1016/j.aquatox.2019.105325

Fitzgerald, J. (2016). The effects of hypoxia on chemical toxicity in two model fish species (dissertation).

Friedrich, J., Janssen, F., Aleynik, D., Bange, H. W., Boltacheva, N., Çagatay, M. N., Dale, A. W., Etiope, G., Erdem, Z., Geraga, M., Gilli, A., Gomoiu, M. T., Hall, P. O., Hansson, D., He, Y., Holtappels, M., Kirf, M. K., Kononets, M., Konovalov, S., … Wenzhöfer, F. (2014). Investigating hypoxia in aquatic environments: Diverse approaches to addressing a complex phenomenon. Biogeosciences, 11(4), 1215–1259. 10.5194/bg-11-1215-2014

Government of Canada. (2023, September 12). Government of Canada. Canada.ca. https://www.canada.ca/en/campaign/critical-minerals-in-canada/canadian-critical-minerals-strategy.html

Government of Canada. (2024, March 15). Rare earth elements facts. Natural Resources Canada. https://natural-resources.canada.ca/our-natural-resources/minerals-mining/mining-data-statistics-and-analysis/minerals-metals-facts/rare-earth-elements-facts/20522

Jin, H., Afiuny, P., Dove, S., Furlan, G., Zakotnik, M., Yih, Y., & Sutherland, J. W. (2018). Life cycle assessment of neodymium-iron-boron magnet-to-magnet recycling for Electric Vehicle Motors. Environmental Science &amp; Technology, 52(6), 3796–3802. 10.1021/acs.est.7b05442

Liu, S.-L., Fan, H.-R., Liu, X., Meng, J., Butcher, A. R., Yann, L., Yang, K.-F., & Li, X.-C. (2023). Global Rare Earth Elements Projects: New Developments and supply chains. Ore Geology Reviews, 157, 105428. 10.1016/j.oregeorev.2023.105428

Mandic, M., Best, C., & Perry, S. F. (2020). Loss of hypoxia-inducible factor 1α affects hypoxia tolerance in larval and adult zebrafish (danio rerio). Proceedings of the Royal Society B: Biological Sciences, 287(1927), 20200798. 10.1098/rspb.2020.0798

Mandic, M., Tzaneva, V., Careau, V., & Perry, S. F. (2018). HIF-1α paralogs play a role in the hypoxic ventilatory response of larval and adult zebrafish (danio rerio). Journal of Experimental Biology. 10.1242/jeb.195198

Markich, S. J., Hall, J. P., Dorsman, J. M., & Brown, P. L. (2024). Toxicity of rare earth elements (rees) to marine organisms: Using species sensitivity distributions to establish water quality guidelines for protecting Marine Life. Environmental Research, 261, 119708. 10.1016/j.envres.2024.119708

Milsom, W. K. (2012). New insights into Gill chemoreception: Receptor distribution and roles in water and air breathing fish. Respiratory Physiology &amp; Neurobiology, 184(3), 326–339. 10.1016/j.resp.2012.07.013

Neubauer, P., & Andersen, K. H. (2019). Thermal performance of fish is explained by an interplay between physiology, behaviour and ecology. Conservation Physiology, 7(1). 10.1093/conphys/coz025

Paquin, P. R., Gorsuch, J. W., Apte, S., Batley, G. E., Bowles, K. C., Campbell, P. G. C., Delos, C. G., Di Toro, D. M., Dwyer, R. L., Galvez, F., Gensemer, R. W., Goss, G. G., Hogstrand, C., Janssen, C. R., McGeer, J. C., Naddy, R. B., Playle, R. C., Santore, R. C., Schneider, U., … Wu, K. B. (2002). The biotic ligand model: A historical overview. Comparative Biochemistry and Physiology Part C: Toxicology &amp; Pharmacology, 133(1–2), 3–35. 10.1016/s1532-0456(02)00112-6

Perry, S. F., Jonz, M. G., & Gilmour, K. M. (2009). Chapter 5 oxygen sensing and the hypoxic ventilatory response. Fish Physiology, 193–253. 10.1016/s1546-5098(08)00005-8

Philippe, C., Vergauwen, L., Huyghe, K., De Boeck, G., & Knapen, D. (2023). Chronic handling stress in zebrafish danio rerio husbandry. Journal of Fish Biology, 103(2), 367–377. 10.1111/jfb.15453

Playle, R. C. (1998). Modelling metal interactions at fish gills. Science of The Total Environment, 219(2–3), 147–163. 10.1016/s0048-9697(98)00232-0

Prabhakar, N. R., & Semenza, G. L. (2012). Adaptive and maladaptive cardiorespiratory responses to continuous and intermittent hypoxia mediated by hypoxia-inducible factors 1 and 2. Physiological Reviews, 92(3), 967–1003. 10.1152/physrev.00030.2011

Santore, R. C., Ryan, A. C., Kroglund, F., Rodriguez, P. H., Stubblefield, W. A., Cardwell, A. S., Adams, W. J., & Nordheim, E. (2017). Development and application of a biotic ligand model for predicting the chronic toxicity of dissolved and precipitated aluminum to aquatic organisms. Environmental Toxicology and Chemistry, 37(1), 70–79. 10.1002/etc.4020

Small, K., Kopf, R. K., Watts, R. J., & Howitt, J. (2014). Hypoxia, Blackwater and fish kills: Experimental lethal oxygen thresholds in juvenile predatory lowland river fishes. PLoS ONE, 9(4). 10.1371/journal.pone.0094524

Tellier, J. M., Kalejs, N. I., Leonhardt, B. S., Cannon, D., Höök, T. O., & Collingsworth, P. D. (2022). Widespread prevalence of hypoxia and the classification of hypoxic conditions in the Laurentian Great Lakes. Journal of Great Lakes Research, 48(1), 13–23. 10.1016/j.jglr.2021.11.004

Vukov, O., Smith, D. S., & McGeer, J. C. (2016). Acute dysprosium toxicity to Daphnia Pulex and Hyalella Azteca and development of the Biotic Ligand Approach. Aquatic Toxicology, 170, 142–151. 10.1016/j.aquatox.2015.10.016

Zachar, P. C., & Jonz, M. G. (2012). Neuroepithelial cells of the gill and their role in Oxygen Sensing. Respiratory Physiology &amp; Neurobiology, 184(3), 301–308. 10.1016/j.resp.2012.06.024

